# Does pollen limitation limit plant ranges? Evidence and implications

**DOI:** 10.1101/2021.08.18.456861

**Authors:** Emma Dawson-Glass, Anna L. Hargreaves

## Abstract

Range limits often involve declines in sexual reproduction, reducing fitness, dispersal, and adaptive potential at range edges. For plants, sexual reproduction is frequently limited by inadequate pollination. While case studies show that pollen limitation can limit plant distributions, the extent to which pollination commonly declines toward plant range edges is unknown. Here, we leverage global databases of pollen-supplementation experiments and plant occurrence data to test whether pollen limitation increases toward plant range edges, using a phylogenetically controlled meta-analysis. While there was significant pollen limitation across studies, we found little evidence that pollen limitation increases toward plant range edges. Pollen limitation was not stronger toward the tropics, nor at species’ equatorward vs poleward range limits. Meta-analysis results are consistent with results from targeted experiments, in which pollen limitation increased significantly toward only 14% of 14 plant range edges, suggesting that pollination contributes to range limits less often than do other interactions. Together, these results suggest pollination is one of the rich variety of potential ecological factors that can contribute to range limits, rather than a generally important constraint on plant distributions.

## Introduction

All species have limited geographic ranges, and the causes of these range limits shape ecological communities and species responses to global change. In the short term, species ranges are limited ecologically by environmental constraints on fitness, and/or species’ ability to disperse to suitable habitat beyond their current range (1). In the longer term, ranges can be limited evolutionarily by lack of ability to adapt their way out of fitness and dispersal limitations. A critical life stage for all three constraints (fitness, dispersal, adaptation) is sexual reproduction, which often declines toward or beyond range edges (2; 3). Lower sexual reproduction at range edges reduces individual fitness, reduces the number of offspring/propagules that can disperse to and potentially colonize beyond range habitat, and erodes the genetic diversity needed to adapt to beyond-range conditions by reducing recombination and gene flow (4). These effects are particularly pronounced for plants, as pollen and seeds (the precursor to and product of sexual reproduction, respectively) are generally the only opportunity for long-distance geneflow and dispersal. Thus sexual reproduction is particularly important in the evolutionary ecology of plant range limits.

Sexual reproduction can decline toward range limits due to lack of resources and/or lack of mates (or for plants, lack of pollen). Many range limits occur along ecological gradients, where the *quality* of a species’ habitat declines until populations are no longer self-sustaining (2). As reproduction is energetically costly, declining habitat quality can reduce reproduction directly via lack of resources. Range limits may also arise due to declines in the *quantity* of suitable habitat (5). If declines in habitat quality or quantity lead to smaller or sparser populations toward range edges (3), sexual reproduction can be limited by lack of compatible mates (6). For example, plants in small range-edge populations of a wind-pollinated grass received fewer pollen grains than plants in dense central populations, and so made fewer seeds (7). For animal-pollinated plants, reproduction can also be limited by lack of pollinators. Pollinator limitation can arise if pollinators become scarce along an abiotic gradient (8, 9), or if small plant populations or floral displays fail to attract consistent pollinators (10). Thus, classic theory that individual fitness, population size, and population density decline toward range limits (11) suggests several mechanisms by which reproduction might become increasingly pollen limited toward plant range edges.

Alternatively, pollen limitation need not increase toward plant range edges. While the size of populations and individuals often declines toward range edges, this pattern is not universal (3), and if edge populations do not have smaller floral displays, they should not be disadvantaged in attracting consistent pollinator visitation. Even when edge populations do attract fewer pollinators, plants may be able to compensate. Some alpine plants can offset low daily visitation by increased flower lifespan (12). Chronic pollen limitation can lead to adaptations for increased self-pollination, including higher self-compatibility and autonomous pollen deposition (i.e. without the need for a pollen vector) (13; 14). Indeed, if populations frequently become more selfing toward range edges (15), we might find the opposite pattern, whereby pollen limitation could decline toward range edges (note that increased selfing would reduce only the ecological effects of pollen limitation, not the evolutionary effects of reduced geneflow). Finally, if edge populations suffer from lack of resources, seed set may ultimately be resource limited even if pollen receipt also declines–in other words, plants would not be able to produce more seeds even if they had more high-quality pollen (16).

Whether seed production is limited by resources or pollination has interesting ecological and evolutionary consequences for plant populations, and so has been the subject of much study (e.g. 17–19). Biologists typically measure pollen limitation by experimentally adding outcross pollen to some stigmas. If pollen-supplemented flowers produce more seeds than un-supplemented controls, seed set is limited by the quantity or quality of pollen receipt (20). Pollen-supplementation experiments are often done at a single site, but multi-site studies and syntheses have found geographic patterns. Pollen limitation tends to be higher toward low latitudes, apparently driven by increasing plant diversity (21) and shows varied patterns with elevation depending on the region (16, 22). Despite a 20+ year history of comparing pollen limitation among populations and long-standing theory that pollen limitation should increase toward plant range edges, few studies have assessed whether insufficient pollination constrains plant distributions (23) and only a dozen or so have explicitly tested whether pollen limitation increases toward plant range edges (Table 1).

**Table 1.** Studies that explicitly tested whether pollen limitation increases toward plant range edges. Compiled by searching the references and citing literature in each study in the table, starting with the earliest five tests of the hypothesis (i.e. 2015 or earlier).

| Plant species | Reliance on pollinators | Populations surveyed | Type of range limit | Change in pollen limitation toward range edge? | Study |
| --- | --- | --- | --- | --- | --- |
| *Rhinanthus minor*<br>(Orobanchaceae) | Self-compatible,<br>capable of full<br>autonomous pollination | 6: two transects from range<br>core to edge | High elevation | No change<br>(no populations pollen<br>limited) | Hargreaves et<br>al 2015 [14] |
| *Campanula*<br>*americana*<br>(Campanulaceae) | Self-compatible,<br>capable of limited<br>autonomous pollination | 24: spanning range | Low latitude<br>High latitude | No change<br>(most populations pollen<br>limited) | Koski et al<br>2017 [26] |
| *Erythronium*<br>*montanum*<br>(Liliaceae) | Self-compatible,<br>capable of limited<br>autonomous pollination | 4: transect from low edge<br>to high edge | Low elevation<br>High elevation | Non-significant increase | Theobald et al<br>2016 [27] |
| *Clarkia xantiana* ssp<br>*xantiana* **<br>(Onagraceae) | Self-compatible,<br>not capable of<br>autonomous pollination | 12: transect from range<br>core to edge | Low elevation /<br>dry | Increase | Moeller et al<br>2012 [9] |
|  |  | 6: transect from range core<br>to beyond range*<br>3: range core, range edge,<br>beyond range* |  | No change<br>(but stronger beyond range<br>than in range) | Benning &<br>Moeller 2019<br>[28]<br>Anderson et al<br>2015 [29] |
| *Clarkia xantiana* ssp<br>*parviflora* **<br>(Onagraceae) | Self-compatible | 3: range core, range edge,<br>beyond range* | High elevation /<br>wet | No change<br>(but stronger beyond range<br>than in range) | Anderson et al<br>2015 [29] |
| *Erythronium*<br>*americanum*<br>(Liliaceae) | Partially self-incompatible | 6: two transects of 3 sites<br>from range core to edge | High elevation | No change | Rivest &<br>Vellend 2018<br>[30] |
| *Trilium erectum*<br>(Melanthiaceae) | Partially self-incompatible | 6: two transects of 3 sites from range core to edge | High elevation | Increase | Rivest & Vellend 2018 [30] |
| *Witheringia solanacea*<br>(Solanaceae) | Self-incompatible, | 4: two pairs of range core and range edge | High elevation | No change | Stone & Jenkins 2008 [15] |
| *Astragalus utahensis*<br>(Fabaceae) | Self-incompatible | 8: four core, four edge | High latitude | No change | Baer & Maron 2018 [31] |
| *Lilium pomponium*<br>(Liliaceae) | Self-incompatible | 17: throughout range | Low latitude<br>High latitude | Non-significant increase<br>No change | Macri et al 2021 [32] |
| *Embothrium coccineum*<br>(Proteaceae) | Self-incompatible | 16: rough transect from centre to edge | Low elevation / increased aridity | Increase | Chalcoff et al 2012 [8] |

\*pollen limitation assessed on transplanted populations so could be measured beyond range
\*\*species excluded from meta-analysis as subspecies range limits were in the core of the overall species range; ie sites closest to each subspecies range centre would appear to be closest to the range edge and vice versa

To overcome the lack of targeted studies measuring pollen limitation toward plant range edges, we combine two large, public data sets. We accessed pollen-limitation estimates from the GloPL database, which compiles the results of pollen supplementation experiments (24). For every species with a pollen-limitation estimate in GloPL, we looked for occurrence data on the Global Biodiversity Information Facility (GBIF; 25) from which we could generate a species distribution polygon. We then calculated how close each pollen supplementation experiment was to the species’ nearest range edge. If pollen limitation is generally an important contributor to plant range limits, we predict it will increase as distance to the range edge decreases, both within species for species with pollen-limitation estimates from >1 site (multi-experiment data), and among species (full data).

We also test three additional factors that may influence geographic patterns in pollen limitation. First, given previous findings that pollen limitation increases toward the tropics (21), we include a covariate for absolute latitude. Second, long-standing speculation posits that biotic interactions are more likely to limit the low-latitude ends of species distributions (11), with strong empirical support at least for negative interactions (23). We therefore include a covariate that indicates whether the nearest range edge is toward the polar or equatorward end of the species latitudinal distribution. Finally, the expectation that pollen limitation increases at range edges rests largely on the assumption that fitness or population size/density decline toward range limits, which in turn assumes that range edges occur across continuous environmental gradients, rather than being stopped abruptly by barriers like oceans. We therefore include a categorical predictor indicating whether the nearest range edge is imposed by an ocean or occurs across continuous land, and test whether the effect of distance from the range edge varies between these two range limit types.

## Methods

All data manipulation and analyses were done in R (R Core Team, 2020); all code will be publicly archived upon acceptance.

### Pollen limitation data (GloPL)

We obtained estimates of pollen limitation from the GloPL database, which contains results from 2969 pollen-supplementation experiments compiled from >900 publications on >1200 wild plant taxa (downloaded April 2020; 24). For each study x taxon x replicate (e.g. different years or populations), GloPL provides a pollen-limitation effect size, calculated as the log response ratio of the reproductive output of pollen-supplemented flowers compared to unsupplemented controls:

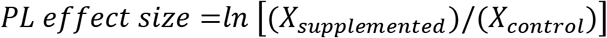

where *X* is the mean reproductive output (24). Thus if pollen supplementation had no effect the effect size is ln(1)=0; if it doubled reproductive success the effect size is ln(2)=0.69, etc. In theory, supplemented flowers should make at least as many seeds as unsupplemented flowers, but due to random variation and potentially damage during supplementation (33) negative effect sizes exist. We generally retained negative effect sizes, but did explore and remove two extreme outliers (SI.1). Studies used various measures of reproductive output; GloPL calculates effect sizes from seeds/plant if available, otherwise gives preference to responses in the order: seeds/flower, seeds/fruit, fruits/flower, seeds/ovule. As log transformations cannot handle zeros, GloPL added 0.5 to zero values (2.4% of response ratios) before transformation (24).

Meta-analyses require an estimate of sampling variance, which we calculated for each GloPL effect sizes as:

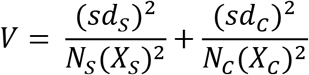

where subscripts *S* and *C* refer to the pollen-supplemented and unsupplemented-control treatments respectively, *sd* is the standard deviation, and *N* is the number of replicates (34). We excluded 181 effect sizes for which we could not calculate sampling variance (e.g. missing *sd* or *N*), which excluded 81 species. Because zero variances cause convergence failure of statistical models, we added a small constant (0.00001) to every sampling variance in our data, as per (35).

Finally, we downloaded the phylogenetic tree created for species in GloPL (24) and extracted the geographic coordinates for each pollen supplementation experiment (24). 18 GloPL experiment locations appeared to be in oceans. We checked these locations in the original studies; if possible, we corrected the coordinates, otherwise we discarded these effect sizes (2 discarded).

### Species distributions (GBIF)

For every species with a pollen limitation effect size, associated variance and location, we searched for species-level occurrence data on GBIF (25), downloaded between May and Aug 2020. Occurrence data were cleaned as follows. We used the `CoordinateCleaner` package (36) to remove records: in oceans; with identical latitude and longitude; within a 1° radius of GBIF headquarters, 100 m radius of known biodiversity institutions, or 1 km radius of country centroids or administrative capitals; based on fossil, machine, or living specimen collections (e.g. in a botanical garden); collected before 1945; that were impossible (e.g. latitude >90); and repeat occurrences.

We had to decide how to handle islands, as species distributions and pollinator communities are notoriously affected by island biogeography (37), and because it was often difficult to determine range edges across fragmented archipelagos. We decided to exclude occurrences on islands >200 km from mainland (e.g. New Zealand, Hawaii) and highly fragmented archipelagoes (e.g. Indonesia). Islands with large contiguous areas and good GBIF coverage (e.g. Japan) were retained.

From occurrence data, we created polygons to serve as range maps. We first mapped each species’ occurrence records and GloPL experiment locations, to assess which species had sufficient data to generate a polygon. For species that occur in multiple non-contiguous continents, we retained only points in continents with GloPL experiments (if >1 continent had GloPL experiments we made one range polygon per continent). We generated a concave polygon around external occurrence points with 2 degrees concavity (`concaveman` package; 38), and removed with any polygon area that overlapped oceans. We chose a concave structure as convex hulls often fail to capture nuance and concavity in true distributions (39), and chose 2 degrees concavity as it provided the finest resolution of edge structure without becoming so angular that polygons frequently intersected themselves and ceased to be polygons. We excluded 124 GloPL species with <3 GBIF occurrence records or whose polygons were self-intersecting.

We evaluated whether each GBIF-derived range polygon seemed plausible and reliable for determining the distance between pollen-supplementation experiments and the nearest range edge. We considered polygons implausible if GloPL experiments were far outside them or if polygons stopped at country borders, and excluded these species. For 75 GloPL experiments that were outside but within ~10 km of the polygon, we redrew the polygon including the GloPL experiment location as an occurrence (main results), and explored their effect by re-running analyses without them (SI.2). When polygons were very geographically restricted, derived from few occurrence points, or visually unusual (e.g. had extreme outliers, distinct clusters within a continent, wide ranging but extremely patchy distributions), we tried to verify the species range using independent descriptions from national online floras (e.g. Flora of North America), or species-specific literature. If this confirmed outlier points were errors, we manually excluded them. If it confirmed species had distinct occurrence clusters separated by large expanses of unoccupied space, we considered the range limits of the cluster with the GloPL experiment to be most ecologically relevant and removed other clusters. If we could not verify a species’ range or reliably determine a range edge (e.g. species widely and patchily distributed) we excluded it from analyses (Fig. S2). At the end of this data cleaning, we had both pollen-limitation data and range polygons for 563 species x continent combinations.

### Data extraction from range maps

#### Distance to range edge

We calculated a proportional measure of the distance from each pollen supplementation experiment to the nearest range limit (`geosphere` package, 40). We used proportional distance as the biological meaning of a linear distance will vary depending on species’ dispersal ability and the structure of the range edge (41). For each species we identified the centroid of its range polygon, and for each GloPL experiment we identified the nearest point on the species’ range edge (‘edge point’; Fig. 1). We then measured the distances from the experiment to the edge point and to the range centroid, accounting for Earth’s curvature. We calculated the proportional distance to the range edge as:

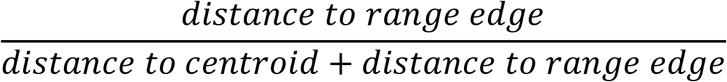

**Fig 1.**
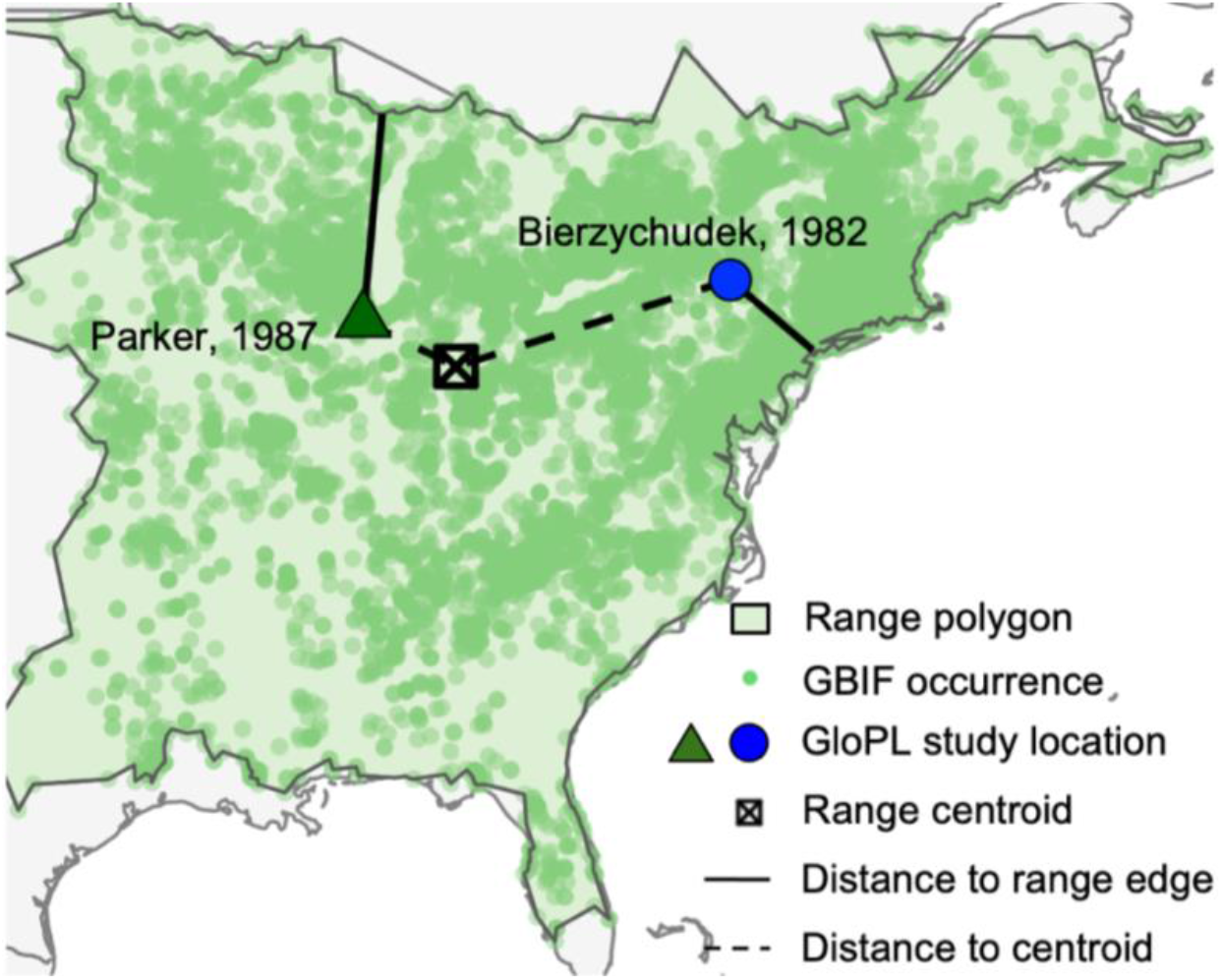
Calculating distance to nearest range edge. Example shows pollen supplementation experiments (studies 61, 62 in GloPL; large points), and GBIF occurrence data (small green circles) for *Arisaema triphyllum*.

Because there is more area close to the edge of a polygon than close to the centre, the distribution of proportional distance measures clusters toward the range edge; however this clustering is less extreme than if we used raw distance (Fig. S3).

#### Edge type

We visually assessed whether the nearest range edge for each pollen supplementation experiment occurred over continuous land or was imposed by an ocean (using the maps above), generating a binary variable (‘land’ or ‘ocean’).

#### Polarity

As biotic interactions are often proposed to be more limiting at the low-latitude ends of species ranges, we tested whether pollen limitation increases more toward species equatorward vs poleward range edges. To do so, we created a measure of ‘polarity’, which quantifies whether a point on a given range edge is toward the polar (*Polarity*=0) or equatorward (*Polarity*=180) end of the species range. We calculated the cardinal direction from the GloPL experiment to the nearest edge point. For the Northern hemisphere, this yields bearing=0 for a polar edge, 180 for an equatorward edge, 90 and −90 for East and West, respectively. For the Southern hemisphere, this yields the reverse (e.g. polar=180); to correct this we subtracted the absolute value of the bearing from 180. We then took the absolute value of all bearing measurements (i.e. both East and West=90).

### Dataset descriptions

We created two datasets (Fig. 2). The first includes only species for which pollen-limitation was assessed at multiple experimental locations (‘multi-experiment dataset’). This is closest to how one would design an experiment to test whether pollen limitation increases toward range limits (Table 1), and enabled us to test the effect of distance to the range edge within species. The multi-experiment dataset contains 875 pollen supplementation experiments from 196 studies on 127 species. The second, ‘full’ dataset, trades within-species resolution for greater geographic and taxonomic coverage, and includes all species with GloPL pollen-limitation data and adequate GBIF range polygons. The full dataset contains 1605 pollen supplementation experiments from 475 studies on 559 species. We examined effect sizes for publication bias using funnel plots, but there were no obvious gaps in the data that would suggest bias (Fig. S4). All analyses described below were performed on both datasets.

**Fig 2.**
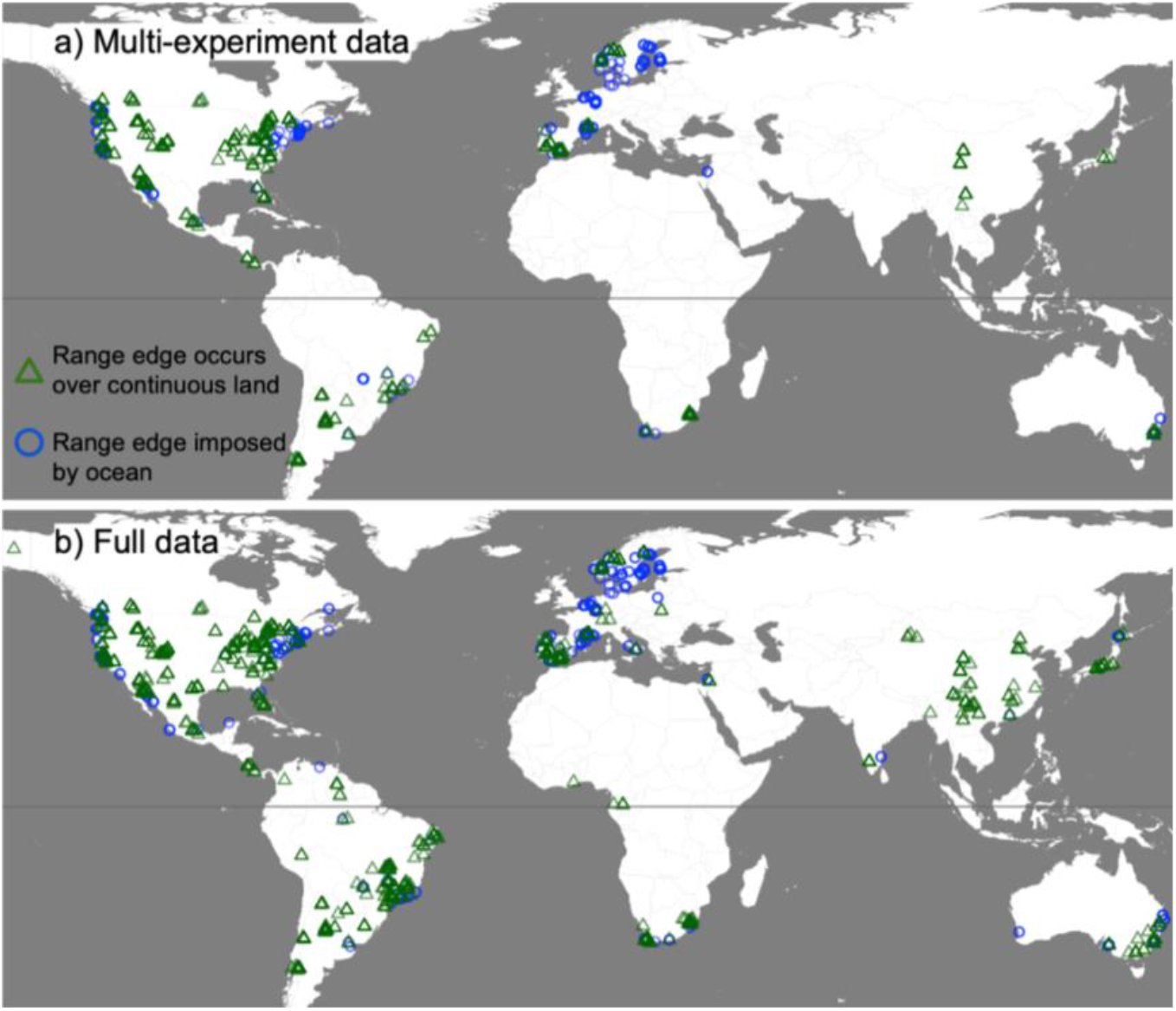
Location of pollen supplementation experiments in our final analyses. a) Multi-experiment data, including only species with pollen supplementation experiments at >1 location. b) Full data.

### Analyses

We conducted a phylogenetically corrected mixed-effects meta-analysis (40) using the ‘rma.mv’ function from the `METAFOR` package (42). The response and fixed effects were:

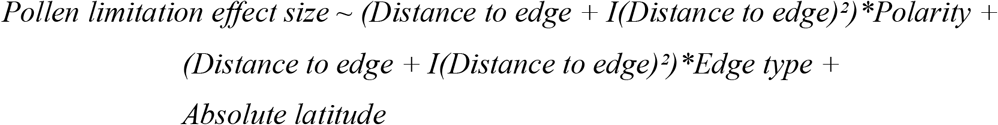

*Pollen limitation effect size* is the effect size from GloPL, and *Distance to edge* quantifies how close the pollen-limitation experiment was to the species range edge (0 at edge to 1 at range centre). We included a quadratic term, *I(Distance to edge)^2^*, as we do not necessarily expect a linear change in pollen limitation from the range core to edge (e.g. pollen limitation might be increase suddenly toward the range edge). The interaction between *Distance to edge* and *Polarity* tests the hypothesis that pollen limitation will increase more strongly toward species’ equatorward range limits. The interaction between *Distance to edge* and *Edge type* tests the hypothesis that pollen limitation will increase more strongly toward range edges that occur across continuous environmental gradients (*Edge type*=land) than those that end abruptly at an ocean (*Edge type*=ocean). *Absolute latitude* tests the hypothesis that pollen limitation is stronger in the tropics, irrespective of where it is measured relative to a species range. We standardized all predictors by centering on 0 and dividing by 2 standard deviations following Grueber et. al. (43). Models weighted effect sizes by the inverse of their sampling variance.

Models included the following random effects. We accounted for phylogenetic distance following (35, 44), by modelling the phylogenetic tree as a correlation matrix (ape package; 45). We included a random intercept for species, to account for heterogeneity due to differences among species unrelated to phylogeny (46). Because experimental methods can affect the estimated magnitude of pollen limitation, we included a random intercept, ‘experimental method’, that denoted all possible combinations of reproductive output measure (seeds/flower, seeds/fruit, etc.), the level at which pollen was supplemented (whole plant, partial plant, or flower), and whether study plants were bagged (47). Finally, to account for non-independence of measurements from the same study and control for overdispersion, we assign a unique ID to each effect size and include a nested random effect for ‘study/effect size’ (46; 48).

To determine the importance of each predictor, we used an information theoretic approach combining model selection and multimodel inference (MuMIn package; 49). We calculated the AICc for our full model and each possible subset of predictors (94 models total). We then created a ‘top set’ of models that were within 2 AICc of the top (i.e. lowest AICc) model (50). For each predictor and interaction present in at least one of the top set of models, we determined its relative influence on pollen limitation by calculating its importance score (sum of the Akaike weights of all top set models that included the parameter), and model weighted-average estimate (i.e. effect size weighted by Akaike weights). We report both measures of influence in our results (Table 2).

**Table 2.** Results from meta-analytical models on pollen limitation. For both datasets, the full model was Pollen limitation effect size ~ (Distance to edge + I(Distance to edge)²) x Polarity + (Distance to edge + I(Distance to edge)²) x Edge type + Absolute latitude. Results are given for parameters in the top set of models. Importance score is the sum of the Akaike weights for each parameter in the top set.

| Dataset | Predictor | Importance score | Model-averaged estimate* & P-value |  |
| --- | --- | --- | --- | --- |
| Multi-experiment data (127 species with pollen supplementation at >1 location) |  |  |  |  |
|  | Intercept |  | 0.589 | **0.010** |
|  | Distance to edge | **0.88** | -0.088 | 0.265 |
|  | Edge type | 0.27 | 0.014 | 0.651 |
|  | Absolute latitude | 0.17 | -0.013 | 0.656 |
|  | Distance to edge x Edge type | 0.14 | 0.044 | 0.607 |
|  | Distance to edge<sup>2</sup> | 0.25 | -0.005 | 0.823 |
| Full data (all 559 species) |  |  |  |  |
|  | Intercept |  | 0.496 | **0.003** |
|  | Distance to edge x Edge type | **1.0** | 0.344 | **0.005** |
|  | Edge type | **1.0** | 0.125 | **0.016** |
|  | Distance to edge | **1.0** | 0.020 | 0.741 |
|  | Absolute latitude | 0.48 | -0.046 | 0.476 |
|  | Polarity | 0.26 | -0.007 | 0.657 |
|  | Distance to edge<sup>2</sup> | 0.21 | 0.006 | 0.746 |

\*predictors standardized by centering and dividing by 2 SD

To visualize results, we made a reduced model including only fixed effects with importance score ≥0.7 (only some of which had significant model-weighted average estimates), in which all parameters were on their original scales, with aariance and random effects as above (51). From this model we generated predicted regression lines and confidence intervals for fixed effects (‘predict’ function; confidence intervals are for visualization only as they do not account for influence of random effects).

## Results

### Multi-experiment data

For 127 species with pollen-limitation effect sizes from multiple experimental locations (875 effect sizes), pollen limitation showed a non-significant increase toward species’ range edges (Fig. 3a). *Distance to edge* was the only term with an importance score >0.7, but its model-weighted average estimate was not significant (*P* > 0.1; Table 2). The intercept was significant, indicating significant pollen limitation detected across studies. The effect of distance to the range edge did not vary between range edges imposed by an ocean vs. those that occurred across continuous land (Table 2). Contrary to biogeographic hypotheses that species interactions are more limiting in the tropics and toward the tropical ends of species ranges, pollen limitation did not vary with absolute latitude, nor was the effect of distance to the range edge stronger for more equatorward range edges (Table 2). Results were consistent if we excluded effect sizes that were originally slightly outside the GBIF-derived range polygons (Table S1; Fig. S1).

**Fig. 3.**
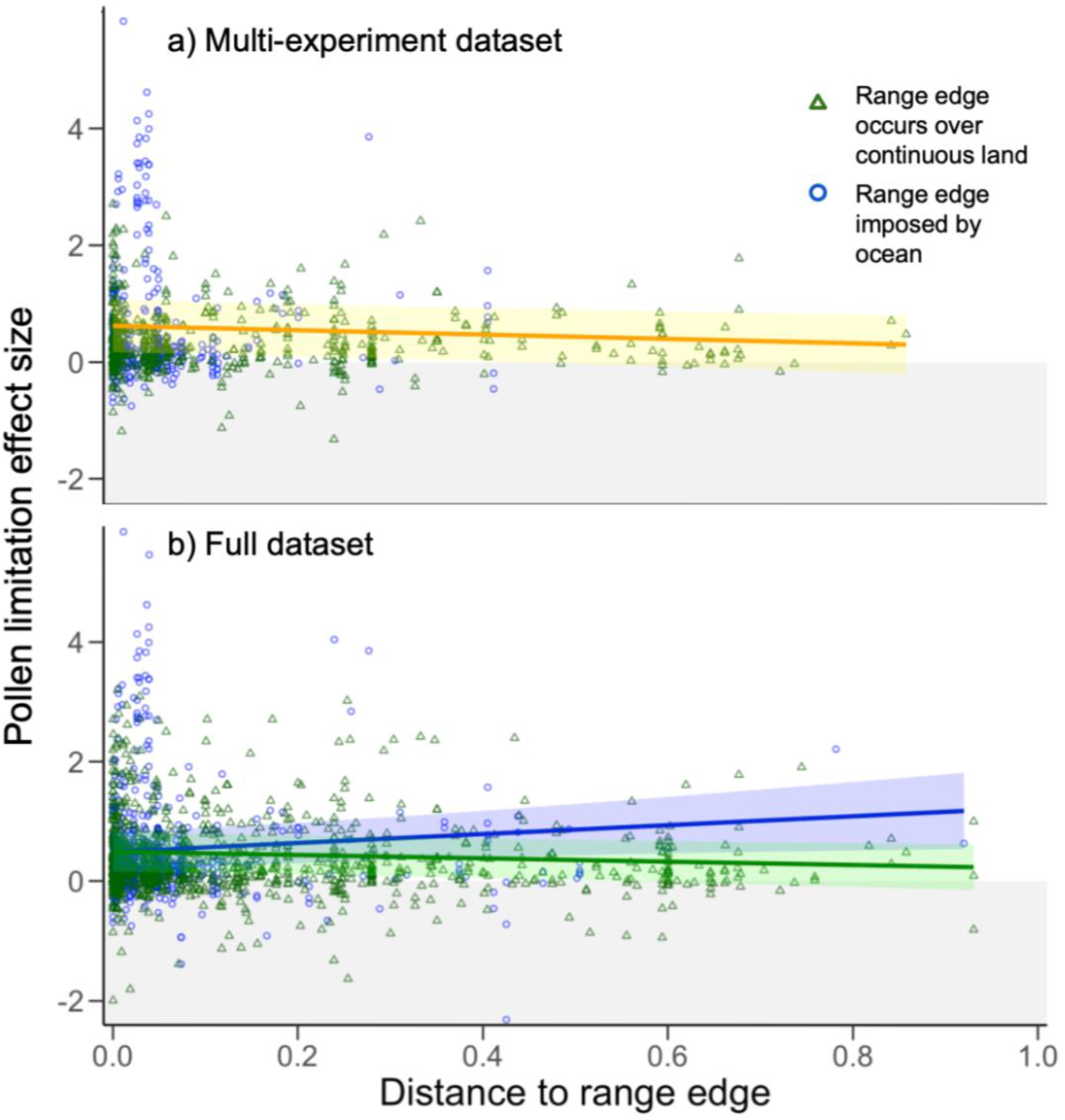
Pollen limitation does not vary strongly toward plant range edges. Points show raw effect sizes: >0 means pollen supplementation increased reproduction; <0 means supplementation decreased reproduction (grey shading indicates this result is theoretically unexpected). Trend lines and 95% confidence intervals are from reduced models that included only parameters with an importance score ≥0.7, not all of which are significant (full statistical results in Table 2). **a)** For 127 species with pollen limitation effect sizes from >1 site, pollen limitation increased non-significantly toward range edges (model: *Pollen limitation ~ Distance to edge*). **b)** For all 560 species, the effect of being near a range edge varied (significantly) between edges imposed by oceans and those occurring across continuous land (model: *Pollen limitation ~ Distance to edge* x *Edge type*).

Whereas 75% of pollen limitation effect sizes ranged from 0 (no pollen limitation) to 2 (pollen supplementation increased reproduction more than 7-fold), there was a cluster of effect sizes >3, meaning supplementation increased reproduction more than 20-fold. We looked more closely at these extremes. Of 19 effect sizes >3 (i.e. the highest 2.2% of effect sizes), 16 came from two species, both of which produce rewardless flowers and so rely on pollination by deceit: *Nerium oleander* (oleander) and *Cypripedium acaule* (pink lady’s slipper orchid). We tested whether these outlier species drove the effect of *Distance to edge* by repeating the analysis without them. Excluding them reduced the importance of *Distance to edge* from 0.88 to 0.67 (as above, no terms had significant model-average estimates; Table S2, Fig S6). Thus, the effect of distance to the range edge was highly sensitive to two of 127 species.

### Full data

When we considered all 560 species (Fig. 2), the effect of distance to the range edge depended on whether the range edge occurred across continuous land or was imposed by an ocean. The *Distance to edge* x *Edge type* interaction appeared in all top models and was significant according to model-averaged estimates (Table 2). However, the form of the interaction was unexpected. Logically, the type of range edge being approached should matter most closest to that range edge. The opposite was true; pollen limitation at land vs. oceanic range edges was essentially identical close to the edge (Fig. 3b). Pollen limitation varied little with range position when the nearest range edge occurred across continuous land; it declined slightly toward edges imposed by oceans, but this effect was driven by higher pollen limitation in the range *core* (Fig. 3b). Thus the *Distance to edge* x *Edge type* interaction says little about processes at play at range edges (and becomes non-significant if we exclude 75 effect sizes originally outside GBIF-derived range polygons; Table S1). Excluding the two species with extreme pollen limitation did not change results (Table S2).

## Discussion

Increasing antagonistic interactions frequently contribute to species range limits, but the role of declining mutualistic interactions is much less understood (23). Given the importance of reproduction to plant range limits, claims that pollination frequently limits plant distributions (52), and newly available data with which to quantify pollen limitation toward range edges (targeted studies and global database of pollen-limitation experiments), it is a natural time to test whether pollen limitation generally intensifies toward plant range edges. Our results suggest that it does not.

Our meta-analytic approach found little support for increasing pollen limitation toward plant range limits. In our biggest dataset (559 plant species), there was no signal that pollen limitation increased toward range edges (Fig. 3b). There is undoubtedly a lot of noise in these data, as pollen limitation was only assessed at one site for 432 of those species and can vary greatly among species (53) even after accounting for phylogenetic relatedness (19). However, when we used a more refined dataset of 127 species for which pollen limitation was measured at multiple sites, pollen limitation increased only slightly and non-significantly toward range limits (Fig. 3a), and even this effect was driven by only two species.

Of course, even in our multi-experiment dataset, few species had pollen-limitation measurements from multiple sites approaching the same range limit; as with many ecological meta-analyses, the data we used were not collected to test the questions we were interested in. We can however compare our meta-analysis results to the growing number of studies that do systematically test pollen limitation toward plant range limits (Table 1). At the 14 range edges where this has been done, there was no change in pollen limitation toward 9, a marginal increase in pollen limitation at 3, and a significant increase at only 2. Together with our meta-analysis results from hundreds of species, this suggests that rather than being a generally important constraint on plant ranges, pollination is but one of the ‘rich panoply’ (54) of ecological factors that limit some range edges but not others.

While we found little evidence that pollen limitation increases toward plant range edges, a few are caveats worth considering. First, the relationship between pollen limitation and resource limitation is notoriously tricky to pin down. For example, one can only supplement pollen when plants flower, yet perennial plants can spend years in non-reproductive stages. If a plant spends 7 years gathering the resources needed to flower and is pollen limited when it finally does, pollen supplementation experiments will find the species is pollen limited, even though 7 years of 8 it is resource limited. Similarly, plants sometimes reallocate resources to reproduction when they receive an unusually high amount or quality of pollen (as is typical with pollen-supplementation), at the expense of reproduction in other flowers or years (55). However, as these examples illustrate, difficulty disentangling pollen- vs resource-limitation generally leads to *over*-estimates of pollen limitation. Thus it seems unlikely such issues are obscuring an otherwise important increase in pollen limitation at range edges.

A second caveat is the inherent variability in both pollen limitation and species range edges. Even for one species at one site, pollen limitation can vary greatly among plants, flowering times, and years (56). This variability makes it hard to detect overall patterns across studies, and meta-analyses of pollen-supplementation experiments often find only weak patterns (19). Species range edges can also be hard to delineate spatially, a problem compounded by our reliance on polygons derived from occurrence data. The location of the polygon edges depends on how concave we make the polygon and the quality of geographic sampling. While we chose reasonable concavity parameters and excluded species with obviously poor sampling, there will inevitably be variation in how accurately our polygons reflect true range edges. Both sources of variation make it more difficult to detect patterns in pollen limitation toward range edges across studies. However, patterns were elusive even among studies that systematically test for pollen limitation toward known range edges (Table 1).

A final caveat is that we could only assess proximity to latitudinal and longitudinal range edges. Plant density and floral displays (14), pollinator abundance (10), and pollen limitation (Table 1) can change steeply toward elevational range edges within species’ wider geographic distributions. Thus there could be important effects of elevational range edges that would be missed by our analyses, and which could add noise that might diminish our ability to detect patterns toward geographic range edges.

### Management and implications

Our results contribute to recent discussions about when to account for biotic interactions when forecasting species distributions. Using Table 1 as a rough guide, pollen limitation contributes to 14% (counting significant results only) to 36% (including non-significant or variable results) of plant range edges. This is considerably lower than some antagonistic interactions; in a recent synthesis of field experiments, competition and predation/herbivory contributed to >50% of range limits at which they were assessed (23). On the other hand, some plants are predictably more likely to suffer from pollen limitation, e.g. obligate outcrossers (19) or those with rewardless flowers (57), and pollinator declines, e.g. species with specialized pollination systems (58). Thus, incorporating pollination might not be a priority relative to other interactions for multi-species modelling efforts, but nevertheless be important for species or clades prone to pollen limitation.

Lack of increased pollen limitation toward range edges may be good news for the evolutionary capacity of edge populations. Pollen limitation is positively associated with selection on traits related to pollination (59), thus more intense pollen limitation toward range limits, had it been common, would have been more likely to impose evolutionary trade-offs that constrain adaptation at range edges (60). Lack of increased pollen limitation toward range edges also suggests that many edge populations are not suffering from reduced gene flow within populations (although does not reveal patterns in geneflow among populations). Thus while pollen limitation is present in many edge populations (Fig. 3), it does not seem to be preventing adaptation at range edges any more than it does in throughout plant distributions.

## Acknowledgements

We thank Joanne Bennett and Jeanne Burns for statistical discussions about analysing GloPL effect sizes, John Benning for discussing *Clarkia* results, Wolfgang Viechtbauer for continual work helping researchers use the metafor package effectively, and the LiberEro foundation for funding for EDG.

